# Structural basis for feedforward control in the PINK1/parkin pathway

**DOI:** 10.1101/2021.08.16.456440

**Authors:** Véronique Sauvé, George Sung, Emma MacDougall, Guennadi Kozlov, Anshu Saran, Rayan Fakih, Edward A. Fon, Kalle Gehring

## Abstract

PINK1 and parkin constitute a mitochondrial quality control system mutated in Parkinson’s disease. PINK1, a kinase, phosphorylates ubiquitin to recruit parkin, an E3 ubiquitin ligase, to mitochondria. PINK1 controls both parkin localization and activity through phosphorylation of both ubiquitin and the ubiquitin-like (Ubl) domain of parkin. Here, we observe that phospho-ubiquitin can bind to two distinct sites on parkin, a high affinity site on RING1 that controls parkin localization, and a low affinity site on RING0 that releases parkin autoinhibition. Surprisingly, NMR titrations and ubiquitin vinyl sulfone assays show that the RING0 site has higher affinity for phospho-ubiquitin than the phosphorylated Ubl. Parkin could be activated by micromolar concentrations of tetra-phospho-ubiquitin chains that mimic a mitochondrion bearing multiple phosphorylated ubiquitins. A chimeric form of parkin with the Ubl domain replaced by ubiquitin was readily activated by PINK1 phosphorylation. In all cases, mutation of the binding site on RING0 abolished parkin activation. The feedforward mechanism of parkin activation confers robustness and rapidity to the PINK1-parkin pathway and likely represents an intermediate step in its evolutionary development.

## INTRODUCTION

Parkinson’s disease is one of the most common neurodegenerative diseases. It causes motor symptoms due to the loss of dopaminergic neurons of the *substantia nigra* in the midbrain. Over 90% of the cases are sporadic and occur late in life. However, 5-10% of the cases are attributed to autosomal mutations that induce the disease in younger patients (Koros *et al*, 2017). Many of these mutations are found in *PARK2* (Kitada *et al*, 1998) and *PARK6* (Valente *et al*, 2004) genes and are responsible for the earliest onset cases. These genes encode respectively for parkin and PINK1 proteins, which work together in a mitochondrial quality control process consisting in tagging proteins of damaged mitochondria with ubiquitin molecules. The accumulation of ubiquitinated proteins at the mitochondrial surface triggers the degradation of either the whole damaged mitochondria through mitophagy (Pickles *et al*, 2018) or the excision of damaged portions through the formation of mitochondrial-derived vesicles (Sugiura *et al*, 2014). Parkin and PINK1 are also involved in the suppression of mitochondrial antigen presentation (Matheoud *et al*, 2016) and activation of the STING pathway (Sliter *et al*, 2018), suggesting a role of inflammation in Parkinson’s disease.

Parkin is a cytosolic E3-ubiquitin ligase composed of a N-terminal ubiquitin-like domain (Ubl) linked to a R0RBR module formed by four zinc-binding domains: RING0, RING1, IBR, and RING2. As an RBR-type E3 enzyme, parkin mediates the transfer of ubiquitin from an E2 enzyme onto a cysteine in the RING2 domain, followed by a second transfer onto the substrate protein (Wenzel *et al*, 2011). X-ray structures of parkin have revealed that it adopts an autoinhibited conformation in basal cell conditions (Riley *et al*, 2013; Trempe *et al*, 2013; Wauer & Komander, 2013). The active cysteine located on the RING2 domain is partially occluded by its RING0 domain., and the E2-binding site on RING1 is blocked by the parkin Ubl domain and a short α-helix referred to as the repressor element of parkin (REP). In cells, parkin needs to both translocate to mitochondria and undergo a conformational change to release the autoinhibition. Parkin recruitment to mitochondria depends on PINK1, a serine/threonine kinase, which acts as a sensor of mitochondrial defects (Geisler *et al*, 2010; Matsuda *et al*, 2010; Narendra *et al*, 2010; Vives-Bauza *et al*, 2010). PINK1 accumulates at the surface of damaged mitochondria when mitochondrial protein import is impaired. There, it phosphorylates ubiquitin molecules present at mitochondrial surface. Parkin binds phosphorylated ubiquitin (pUb) with high affinity, which promotes its accumulation onto mitochondria (Kane *et al*, 2014; Kazlauskaite *et al*, 2014; Koyano *et al*, 2014; Ordureau *et al*, 2014). Parkin is then in turn phosphorylated by PINK1 to activate its ubiquitination activity (Kazlauskaite *et al*, 2015; Kondapalli *et al*, 2012; Kumar *et al*, 2015; Sauve *et al*, 2015; Wauer *et al*, 2015a). The most recent x-ray structures of the complex of phosphorylated parkin (pParkin) and pUb have revealed that the phosphorylation of the parkin Ubl domain (pUbl) is responsible for the structural rearrangement within parkin that exposes the active cysteine (Gladkova *et al*, 2018; Sauve *et al*, 2018). Phosphoserine 65 of pUbl binds a phosphate-binding site on RING0 formed by residues K161, R163 and K211. The relocation of the pUbl domain displaces the catalytic domain RING2 from RING0. The released RING2 domain is free to reposition next to the ubiquitin-charged E2 for the transfer of ubiquitin to the parkin active cysteine and subsequently to target proteins. The addition of new ubiquitin molecules to the mitochondrial outer surface provides more substrates for PINK1, which leads to additional parkin recruitment to mitochondria (Seirafi *et al*, 2015). This positive feedback mechanism amplifies the signal for mitophagy and explains the observation that parkin activity is required for its recruitment (Lazarou *et al*, 2013; Ordureau *et al*, 2014; Shiba-Fukushima *et al*, 2014).

Parkin is also regulated through a feedforward (open-cycle) mechanism which does not require parkin phosphorylation. Multiple studies have shown that deletion of the Ubl domain or loss of serine 65 do not completely abolish parkin recruitment to mitochondria and mitophagy (Ordureau *et al*, 2014; Shiba-Fukushima *et al*, 2012; Tang *et al*, 2017; Zhuang *et al*, 2016). While unphosphorylated parkin can be activated *in vitro* by pUb addition, the molecular mechanism has remained unexplored (Kazlauskaite *et al*, 2014). Here, we confirm the existence of a secondary mechanism for parkin activation that is independent of parkin phosphorylation but dependent on the RING0 pUbl-binding site. We observe that pUb can bind to the pUbl-binding site and, in fact, has higher affinity than the pUbl domain. Experiments with phosphorylated polyUb chains reveal the avidity of the pUb and pUbl-binding sites and mimic the effect of multiple immobilized pUb molecules in proximity to each other on the surface of mitochondria. Finally, experiments with parkin with the Ubl domain replaced by ubiquitin show the chimeric molecule is readily activated by phosphorylation. The feedforward mechanism increases the robustness of the pathway for the clearance of damaged mitochondria and likely represents an early feature in the evolutionary development of the PINK1/parkin pathway.

## RESULTS

### Parkin recruitment and mitophagy in cells

To verify that parkin in cells could be activated in the absence of Ubl phosphorylation, we monitored recruitment of different parkin variants from cytoplasm to mitochondria upon addition of mitochondrial uncoupler carbonyl cyanide *m*-chlorophenyl hydrazone (CCCP) (Figure 1A). WT parkin was actively recruited with over 80% of cells showing puncta of parkin on mitochondria after 2 h (Suppl. Figure S1). The C431S mutant, which has no ligase activity, displayed the lowest recruitment as expected from its incapacity to ubiquitinate mitochondrial proteins and drive the PINK1/parkin feedback cycle. As observed by others, parkin bearing the S65A mutation or deletion of the Ubl domain (ΔUbl parkin) showed partial recruitment even though the mutations prevent parkin phosphorylation. The role of the RING0 pUbl-binding site was evident as recruitment was further reduced by a second mutation (K211N) that disrupts the phospho-serine binding site (Gladkova *et al*, 2018; Sauve *et al*, 2018). These results demonstrate a mechanism of parkin activation that is independent of parkin phosphorylation but dependent on the pUbl-binding site on RING0.

**Figure 1.**
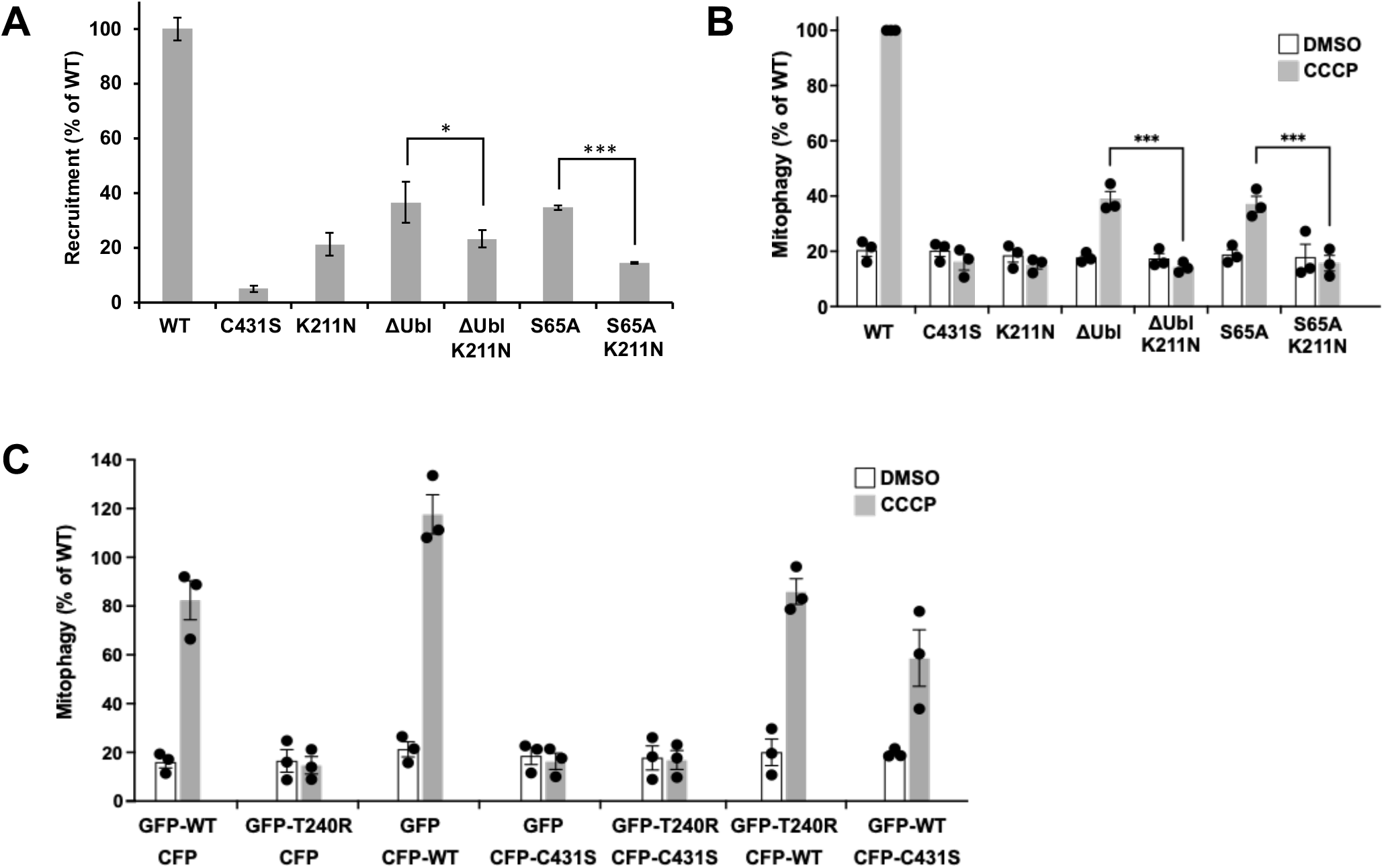
Parkin recruitment and mitophagy without Ubl phosphorylation. **A**. Partial recruitment to mitochondria of parkin mutants without the Ubl-phosphorylation site. Graph shows recruitment of GFP-parkin WT, K211N, C431S, ΔUbl (deletion of residues 1-76), ΔUbl K211N, S65A, S65A K211N measured in U2OS cells 120 min after depolarization of mitochondria by 20 μM CCCP. Mutation K211N of the RING0 pUbl-binding site decreased recruitment even in the absence of Ubl-phosphorylation. Vertical bars represent s.e.m. from two independent experiments. **B**. Mitophagy in the absence of Ubl-phosphorylation. Mitophagy was detected by fluorescence-activated cell sorting (FACS) of untreated and CCCP-treated (20 μM) U2OS cells containing mitochondrially targeted mKeima (mt-Keima) and transiently expressing GFP-parkin. The K211N mutation was epistatic to the Ubl-phosphorylation and decreased mitophagy to the levels seen in the parkin catalytically dead C431S mutant. **C**. Lack of complementation between parkin mutants defective in E2-binding and ubiquitin transfer. U2OS mt-Keima cells were transfected with CFP or GFP alone or as parkin fusion proteins and mitophagy measured by FACS. * P<0.05, **P<0.01, ***P<0.001.

mt-Keima assays were performed to assess the level of mitophagy in cells expressing parkin mutants. The assay measures mitophagy in cells expressing a mitochondrially targeted fluorophore that shifts its excitation spectrum when mitochondria enter the acidic environment of lysosomes. Addition of the protonophore CCCP to depolarize mitochondria induced mitophagy in cells expressing WT parkin (Figure 1B, Suppl. Figure S2). Loss of the parkin active cysteine (C431S) or pUbl-binding site (K211N) abolished the mitophagy response but cells expressing non-phosphorylatable S65A parkin or ΔUbl parkin showed partial mitophagy. The addition of the K211N mutation to inactivate the RING0 pUbl-binding site eliminated the remaining mitophagic activity. These results demonstrate that an alternative mechanism, involving the pUbl-binding site, can trigger mitophagy of depolarized mitochondria.

### Parkin does not show complementation between inactivating mutations

A recurring model of parkin activity has posited that parkin forms dimers or oligomers to allow the catalytic RING2 domain of one molecule to contact the E2∼Ub bound to another molecule (Gundogdu *et al*, 2021). We previously showed an absence of complementation in autoubiquitination assays (Sauve *et al*, 2018) but this left open the possibility of *trans*-active parkin dimers on the surface of mitochondria. To test this, we looked for complementation in the mtKeima assay between the C431S mutant, which is catalytically dead, and the T240R mutant, which is unable to bind E2 enzymes (Figure 1C). The mutants were tagged with GFP and CFP so their expression could be measured and only cells expressing both proteins counted. The mtKeima assay showed robust mitophagy in cells expressing WT parkin after 4 h of CCCP treatment. No mitophagy was detected in cells expressing the T240R and C431S mutants individually or together. Due to variations in the basal signal, the absence of complementation can best be seen in the individual experiments where cells expressing both mutants show no more mitophagy than cells expressing them separately with GFP or CFP (Suppl. Figure S3). These results strongly argue against the existence of *trans-*active parkin oligomers.

### pUb binding activates parkin

We used two different *in vitro* assays to test if pUb binding could activate parkin in the absence of Ubl phosphorylation. The first uses ubiquitin vinyl sulfone (UbVS), which is a chemically reactive derivative that crosslinks to the parkin active site cysteine (Borodovsky *et al*, 2001). The formation of the UbVS adduct is a sensitive measure of exposure of the parkin catalytic site and has been widely used to follow parkin activation (Ordureau *et al*, 2014; Riley *et al*, 2013; Wauer *et al*, 2015a). The advantages of the UbVS assay are its simplicity. It only measures accessibility of the parkin active site cysteine and does not require E1 and E2 enzymes or acyl transfer of ubiquitin to lysine residues. We used the R0RBR construct that is missing the Ubl domain but retains the catalytic RING2 domain and the pUb and pUbl-binding sites on RING1 and RING0. Assays were done with R0RBR parkin purified as a complex with pUb.

UbVS assays showed that addition of pUbl could promote the formation of R0RBR-Ub crosslinks due to release of the RING2 domain from RING0 (Figure 2A). The activation of parkin could be more easily observed with the W403A mutation that partially derepresses parkin activity by destabilizing the REP helix (Trempe *et al*, 2013). Addition of 50 µM pUbl to the W403 mutant generated 50% R0RBR-Ub crosslinks. The high concentration of pUbl required reflects the competition between RING2 and pUbl for binding RING0 and the fact that the RING2 domain is present in the same polypeptide chain at a high local concentration. We used the K211N mutation in the RING0 to confirm the essential role of the pUbl-binding site. No UbVS crosslinks were observed, confirming that release of RING2 was completely dependent on pUbl binding RING0 (Figure 2B).

**Figure 2.**
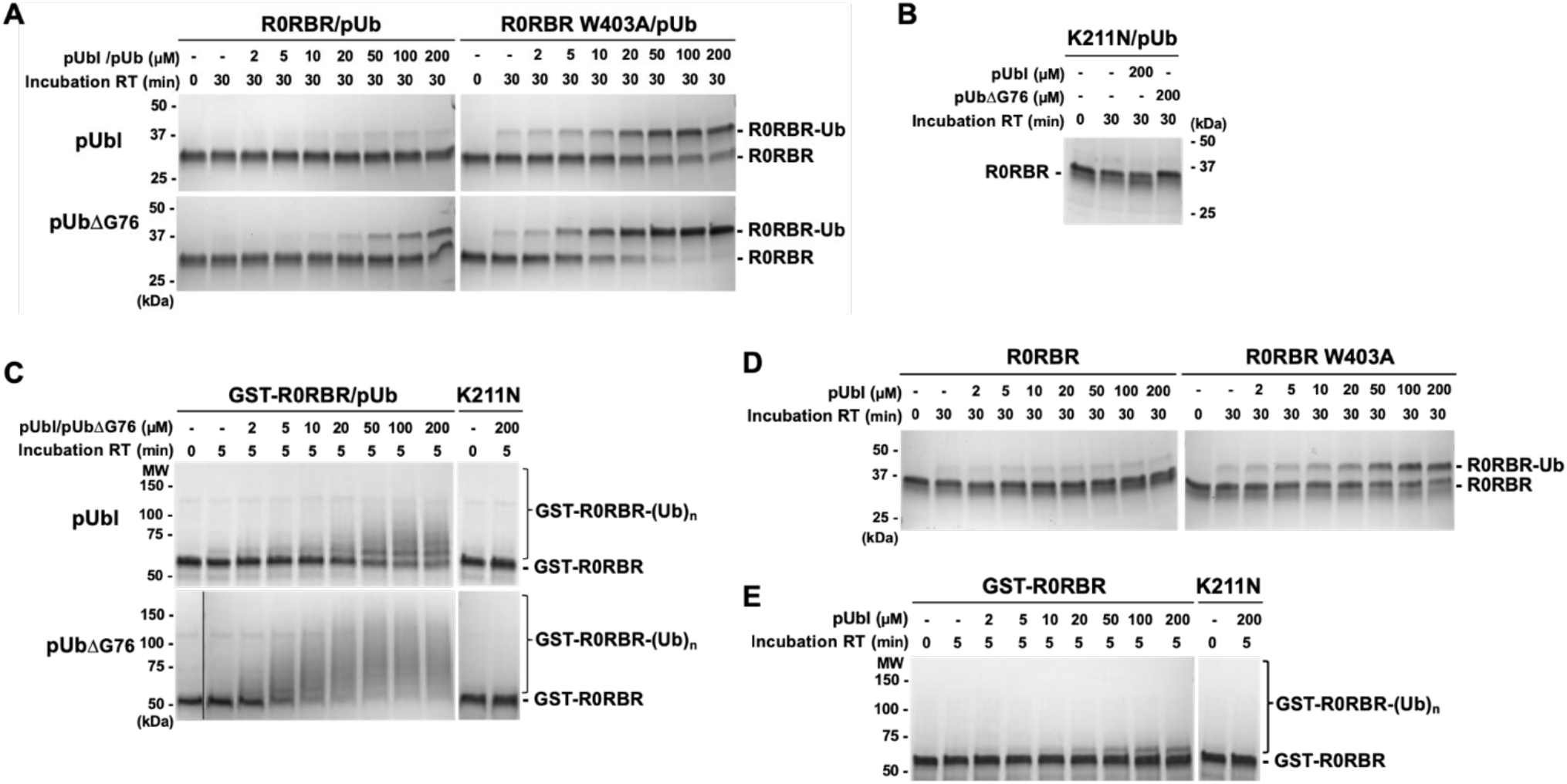
Activation of parkin by pUbl or pUb binding. **A**. Ubiquitin-vinyl sulfone (UbVS) assays of RING2 release by addition of pUbl and pUb *in trans* to R0RBR parkin. Assays were performed with incubation of 2 μM wild-type or W403A R0RBR purified in a 1:1 complex with pUb. Crosslinking was initiated by addition of 10 μM UbVS in the presence of pUbl or pUbΔG76. **C**. Inhibition of R0RBR activation by the K211N mutation. **C**. Autoubiquitination assays of parkin ligase activity induced by pUbl or pUbΔG76 *in trans*. 2 µM GST-R0RBR in complex with pUb was incubated with 50 nM E1, 2 µM UbcH7, 100 µM ubiquitin and 4 mM ATP. **D**. UbVS assays of RING2 release with R0RBR purified without pUb bound. **E**. Autoubiquitination assays of parkin ligase activity with GST-R0RBR without pUb bound.

We next asked if pUb could similarly release the RING2 domain and generate UbVS crosslinks. The assays used pUbΔG76 (pUb without the C-terminal glycine) for consistency with subsequent autoubiquitination assays where pUb interferes with the E2 discharging (Wauer *et al*, 2015b). Similar results were obtained with full-length pUb (Suppl. Figure S4). Surprisingly, pUbΔG76 was much more efficient than pUbl in producing crosslinks, which suggests that it has a higher affinity for the RING0 pUbl-binding site. Addition of pUbΔG76 to wild-type R0RBR led to detectable crosslinking at 20 µM and 50% at 200 µM. The R0RBR W403A mutant was again more sensitive with 50% crosslinking at 7 µM pUbΔG76 and 100% at higher concentrations. Assays with the K211N mutant confirmed that RING0 pUbl-site was required for pUb-induced crosslinks. In contrast, the mutations, H302A, A320R, and N273K, in the RING1 binding sites for pUb and the unphosphorylated Ubl domain had no effect on pUb activation of UbVS crosslinking (Suppl. Figure S4).

In a second set of assays, we measured autoubiquitination of GST-R0RBR parkin upon addition of pUbl or pUb (Figure 2C). In agreement with the UbVS assays, addition of pUbl *in trans* activated parkin ligase activity detected by loss of the unmodified GST-R0RBR parkin and the formation of higher molecular weight bands. pUb was again more efficient than pUbl in releasing parkin autoinhibition and required 10-fold lower concentrations for activation. To confirm the importance of the RING0 Ubl-binding site for parkin activation, we again used a K211N mutant. No autoubiquitination of K211N GST-R0RBR was observed even at 200 µM pUb (Figure 2C).

The UbVS and autoubiquitination assays used R0RBR parkin in complex with a stoichiometric amount of pUb bound to the RING1 site. To investigate possible coupling between the RING1 and RING0 sites, we repeated the pUbl titrations without pUb present. The UbVS assay showed a small decrease in the effectiveness of pUbl additions. W403A R0RBR complexed with pUb required roughly 20 µM pUbl to achieve 50% crosslinking (Figure 2A) while the sample without pUb required 50 µM pUbl (Figure 2D). Autoubiquitination assays showed a much larger difference. Approximately 10-fold more pUbl was required to activate GST-R0RBR parkin in the absence of pUb than in its presence (Figure 2C & 2E).

### Increased local concentration enhances parkin activation by pUb

To compare activation by pUb and pUbl in the context of intact parkin, we designed a chimeric molecule in which the Ubl domain has been replaced by ubiquitin (Figure 3A). To prevent pUb binding to the high affinity site on RING1, we introduced the A320R mutation that disrupts the RING1 site (Wauer *et al*, 2015a). The chimeric protein could be phosphorylated by PINK1 (Figure 3B) and crosslinked to UbVS as efficiently as wild-type parkin (Figure 3C). The RING0 binding site was essential as the K211N mutation prevented crosslinking. Autoubiquitination assays showed that the chimera and wild-type parkin had identical ubiquitination activity when phosphorylated, and activity was again completely blocked by K211N mutation in the pUbl-binding site (Figure 3D). These results demonstrate that ubiquitin can fully replace the Ubl domain of parkin and, when phosphorylated, binds to RING0 to release the catalytic RING2 domain.

**Figure 3.**
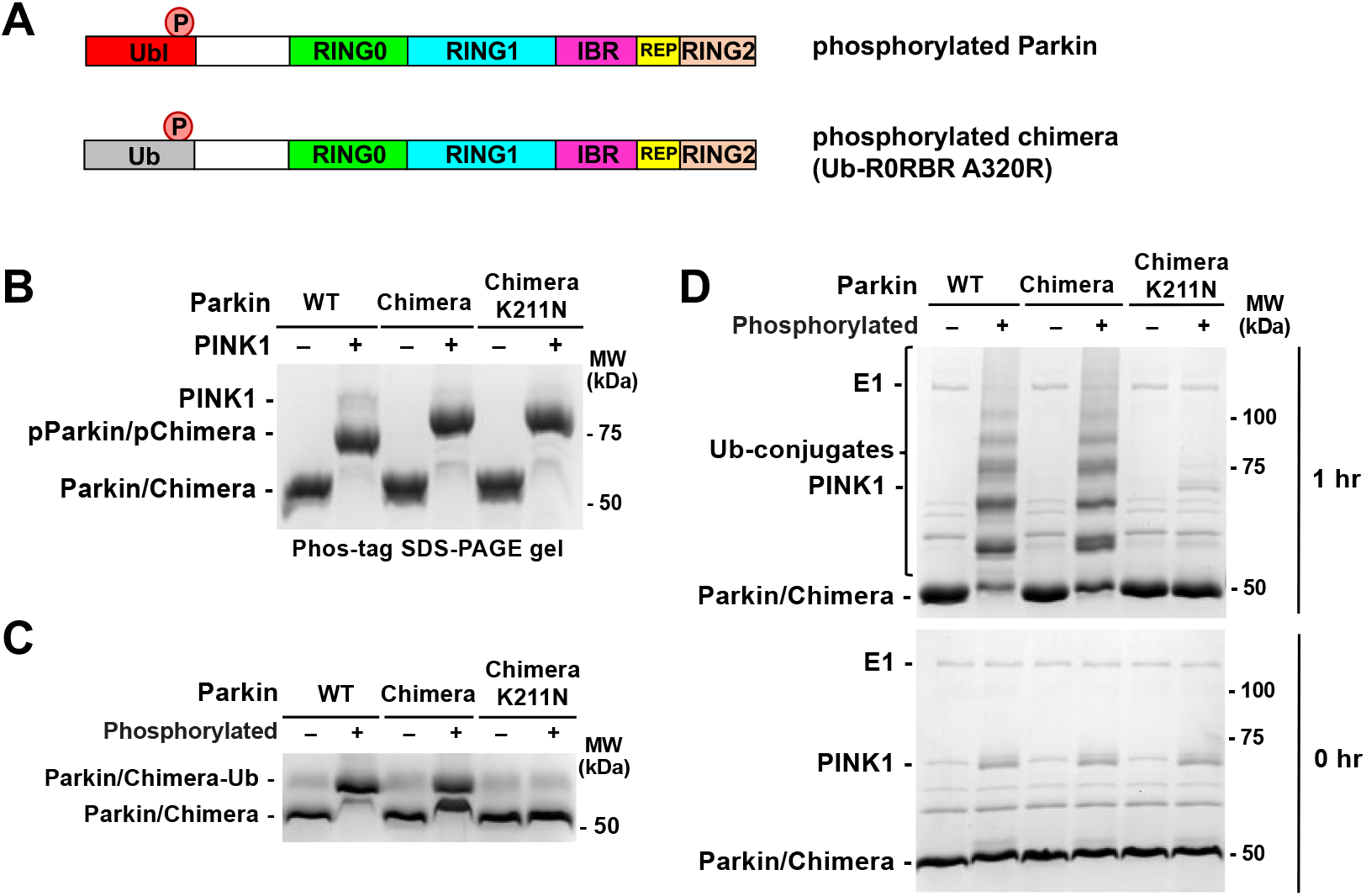
Ubiquitination activity of Ub-R0RBR chimera. **A**. Domains of phosphorylated Parkin (pParkin) and the phosphorylated chimera (pUb-R0RBR) with the A320R mutation. **B**. Phosphorylation of wild-type (WT) parkin and chimeric Ub-R0RBR by PINK1. Reactions were analyzed on a Phos-tag SDS-PAGE gel to assess the level of phosphorylation. **C**. Release of RING2 by Ub-R0RBR chimera. Ubiquitin-vinyl sulfone assays (10 μM UbVS) were performed with 3 μM phosphorylated and non-phosphorylated full-length parkin and Ub-R0RBR. **D**. Ubiquitination activity of Ub-R0RBR chimera. Autoubiquitination assays of phosphorylated full-length parkin and Ub-R0RBR (3.3 µM with 3 µm UbcH7 and 75 µM S65A ubiquitin) were performed to assess the impact of pUbl substitution by pUb on parkin activity.

On damaged mitochondria, PINK1 phosphorylation of ubiquitin molecules likely leads to an elevated local concentration of pUb. We used phosphorylated polyubiquitin chains to mimic the mitochondrial surface modified with multiple pUb molecules. Polyubiquitin chains can be phosphorylated by PINK1 *in vitro* and polyphosphorylated ubiquitin chains have been detected in cells following mitochondrial depolarization (Ordureau *et al*, 2014; Wauer *et al*, 2015b). The existence of two pUb-binding sites on parkin should lead to an increased affinity from cooperation between the sites. Binding of one end of a poly-pUb chain to the high-affinity RING1 site should bring a second pUb molecule in proximity to the low-affinity RING0 site. From modeling based on structures of activated parkin, we determined that the minimal chain length to bridge both pUb binding sites is tetra-pUb (Figure 4A; Suppl. Figure S5).

**Figure 4.**
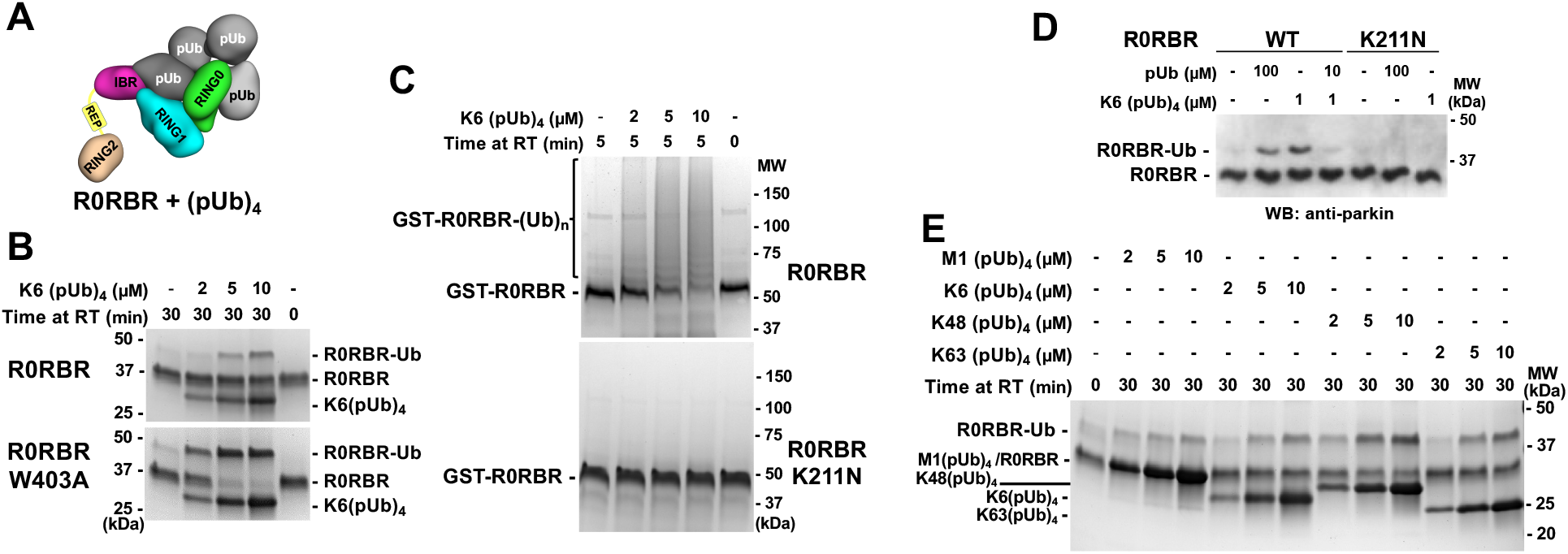
Parkin activation by pUb chains. **A**. Model of (pUb)_4_ chain binding to R0RBR. **B**. UbVS assays of release of RING2 by K6-linked (pUb)_4_ chains. 10 μM UbVS was added to 2 μM R0RBR in the presence of increasing amount of K6(pUb)_4_chains. **C**. Autoubiquitination assays of parkin activation by K6(pUb)_4_. Assays with 2 µM GST-R0RBR, 50 nM E1, 2 µM UbcH7 and 100 µM ubiquitin were performed in the presence of increasing amount of K6(pUb)_4_ chains. **D**. Competition between K6(pUb)_4_ chain and monomeric pUb. UbVs (10 μM) was added to 20 nM of WT and K211N R0RBR in the presence of monomeric pUb and/or K6(pUb)_4_ chain. Reactions were resolved on SDS-PAGE and visualized with anti-parkin antibody by western blotting. **E**. UbVS assays with different types of (pUb)_4_ chains. Assays were performed with 10 μM UbVS and 2 μM R0RBR in the presence of increasing amount of different (pUb)_4_ chains.

We conducted a UbVS assay with K6-linked tetra-pUb chains. Crosslinks were observed at 5 µM K6-linked (pUb)_4_, which is 10-fold lower than with monomeric pUb (Figure 4B). K6(pUb)_4_ chains were also more efficient in crosslinking the W403A mutant although the ability of the assay to measure activation by low concentrations was limited by the need for a stoichiometric amount of pUb chains. We also tested the ability of K6(pUb)_4_ chains to promote the autoubiquitination activity of GST-R0RBR parkin (Figure 4C). Robust autoubiquitination was observed at low concentrations of K6(pUb)_4_ chains and was dependent on the RING0 pUbl-binding site. No ubiquitination activity was observed with the K211N mutant.

To confirm that the activation capacity of tetra-pUb chain was due to binding to RING1 pUb-binding site, we conducted a competition UbVS assay (Figure 4D). We switched to western blot detection which allowed us to use 100-fold less R0RBR (20 nM) in the assay and avoid the stoichiometry issue. The crosslinks observed with 1 µM K6(pUb)_4_ were comparable to those with 100 µM free pUb. When pUb was added at an intermediate concentration (10 µM) that would bind RING1 but not RING0, we observed a reduction of the intensity of the crosslinks induced by 1 µM K6(pUb)_4_. The 10-fold excess of pUb outcompeted K6(pUb)_4_ binding to RING1 and eliminated the cooperativity that promoted K6(pUb)_4_ binding to RING0.

We also tested RING2 release by other commercially available tetra-pUb chains. K48(pUb)_4_ chains were the most efficient in promoting UbVS crosslinking followed by K63(pUb)_4_ and K6(pUb)_4_ chains (Figure 4E). All these chain types are synthesized by parkin and observed *in vivo* (Durcan *et al*, 2014; Ordureau *et al*, 2014). Linear M1(pUb)_4_ chains had a more reduced effect. The levels of activation appear to directly correlate with the ability of the tetra-pUb chains to bridge both pUb and pUbl binding sites. K48(pUb)_4_ chains can easily cover the distance between the two sites and are the least restrained, whereas only one orientation of M1(pUb)_4_ chains can bridge both sites (Suppl. Figure S5).

### Characterization of pUb binding to RING0 by NMR

We previously used NMR spectroscopy to show that pUbl domain could bind R0RBR/pUb complex *in trans* (Sauve *et al*, 2018). Experiments with ^15^N-labeled pUbl showed a loss of signal intensity for phosphoserine 65 upon addition of the complex of R0RBR and pUb. We repeated those experiments with ^15^N-labeled pUb and the R0RBR W403A mutant (Figure 5 & Suppl. Figure S6). Addition of the R0RBR W403A/pUb complex led to selective broadening and loss of signals for pUb residues around the serine 65 phosphorylation site (Figure 5B). When repeated with the double mutant K211N, W403A, the ^15^N-labeled pUb spectrum was unchanged confirming that the changes were the result of pUb binding to the pUbl-site on RING0. When phosphorylated, ubiquitin exists in two conformations due to a shift of a β-strand in the second, minor form (Wauer *et al*, 2015b). We observed similar changes in signal intensity for both conformations implying that both forms bind (Figure 5C). Since the two conformations exchange rapidly in solution (Wauer *et al*, 2015b), measuring the relative affinity of the conformations is difficult, but our results rule out strong specificity for either form.

**Figure 5.**
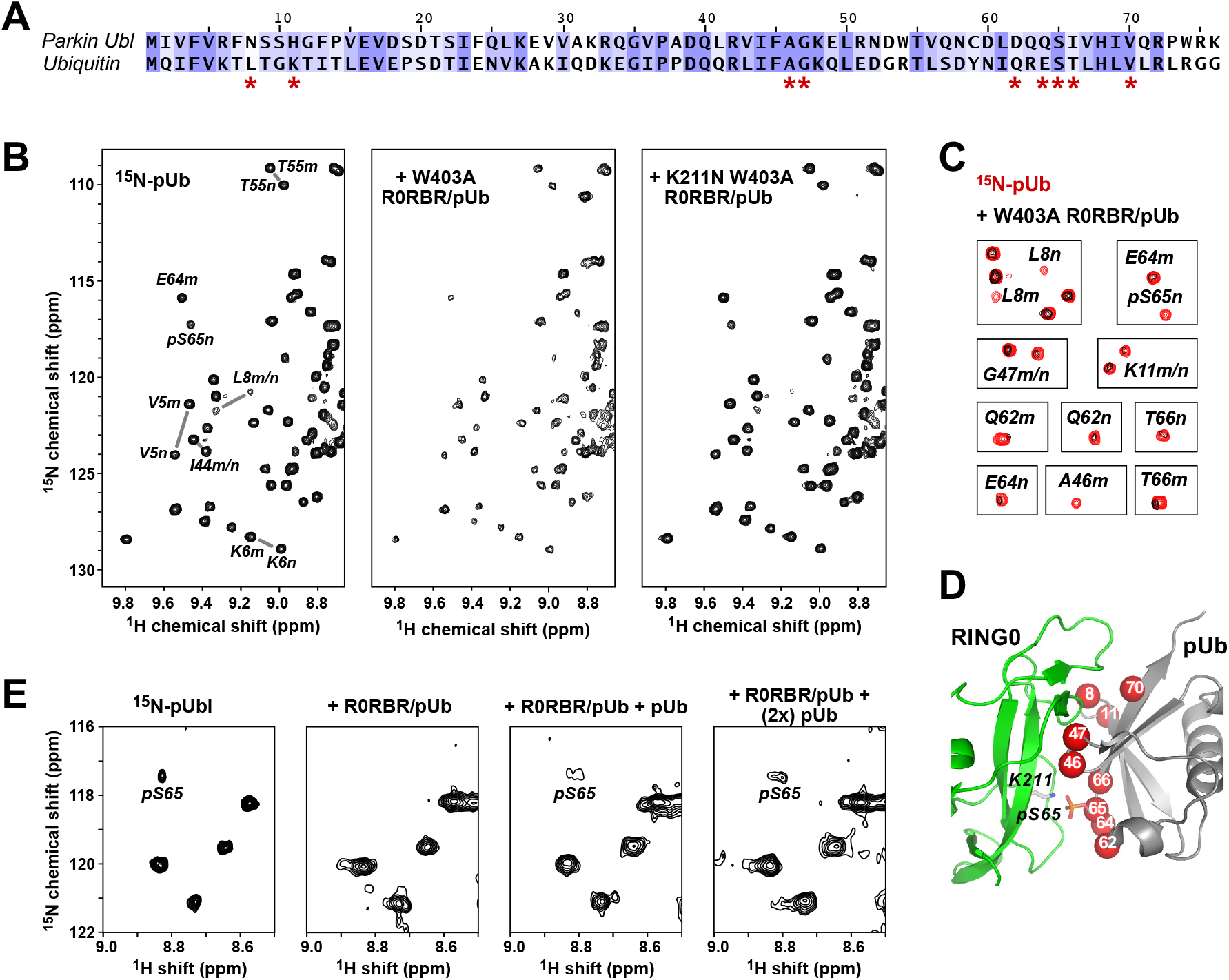
pUb binds to the parkin RING0 domain. **A**. Sequence alignment of ubiquitin and the Ubl domain of human parkin colored according to similarity. Ubiquitin residues whose NMR signals change upon binding RING0 are marked by asterisks. **B**. ^1^H-^15^N correlation spectra at 25 °C of 50 µM ^15^N-pUb alone (*left*) and in the presence of 100 µM complexes of pUb with W403A R0RBR (*middle*) or K211N W403A R0RBR parkin (*right*). Selected backbone amide signals are labeled according to residue number and the pUb conformation, major (m) or minor (n). **C**. Identification of ^15^N-pUb signals that weaken or disappear when pUb is bound to W403A R0RBR. **D**. Mapping of ^15^N-pUb signals that weaken or disappear (*red spheres*) onto a model of pUb bound to parkin RING0. **E**. Competitive binding of pUbl and pUb to RING0. Portion of ^1^H-^15^N NMR correlation spectra at 5 °C of 60 µM ^15^N-pUbl alone (*left*), in the presence of a 1:1 complex of rat parkin R0RBR and pUb (*middle left*), and in the presence of R0RBR/pUb supplemented with 60 μM pUb (*middle right*) and 120 μM pUb (*right*).

We mapped the positions of the residues with the largest changes onto a model of pUb bound to RING0 based on the crystal structures of phosphorylated parkin. Ubiquitin and the parkin Ubl domain share 30% sequence identity and 71% similarity which allows for reliable modelling (Figure 5A). The superimposition of pUb onto pUbl bound to RING0 shows identical conformations except for the loop formed by residues 7-11 (Suppl. Figure S6B). Comparison of the NMR changes with the model shows that the residues with the largest changes all reside at the RING0-pUb interface (Figure 5D). This includes residues leucine 8 and lysine 11 that likely undergo a conformational rearrangement upon binding.

Additional evidence that pUb binds to the parkin pUbl-binding site came from a competition experiment with ^15^N-labeled pUbl (Figure 5E). As previously reported, the pUbl phosphoserine 65 peak disappears when bound to R0RBR/pUb (Sauve *et al*, 2018). Upon addition of excess of pUb, the pS65 peak reappeared confirming the competition for a single binding site.

## DISCUSSION

Previous studies observed the existence of feedforward control in the PINK1/parkin pathway but the molecular mechanism by which pUb directly activates parkin was not identified (Ordureau *et al*, 2014; Shiba-Fukushima *et al*, 2012; Tang *et al*, 2017; Zhuang *et al*, 2016) (Kazlauskaite *et al*, 2014). Here, we show that parkin can be activated without phosphorylation of its Ubl domain and that pUb binding is sufficient to activate ligase activity. pUb binds to the pUbl-binding site allowing pUb to function as a signal for both recruitment of parkin to mitochondria and its activation (Figure 6).

**Figure 6.**
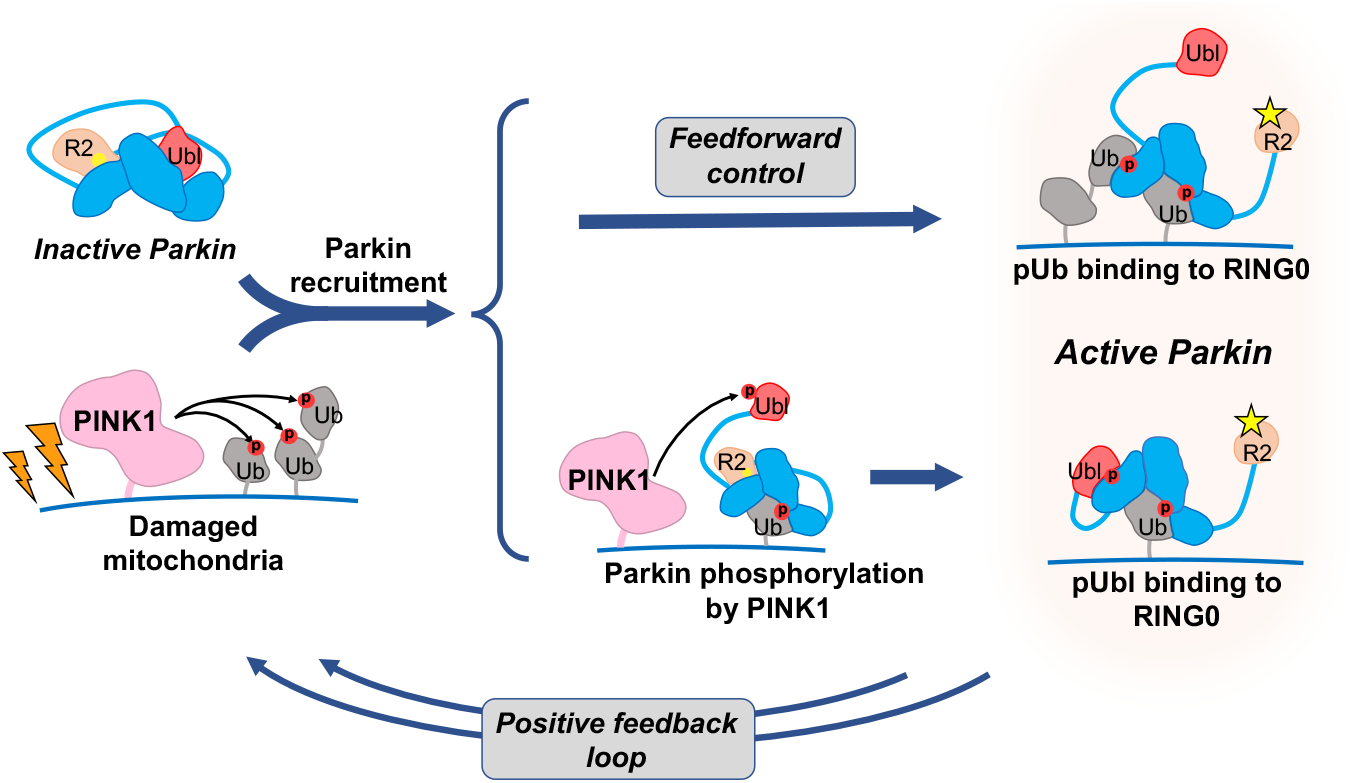
Phosphorylation of ubiquitin on mitochondria acts on both recruitment and activation. Accumulation of PINK1 leads to phosphorylation of ubiquitin on damaged mitochondria and recruitment of inactive parkin. Parkin can be activated through one of two ways. It can bind a second phosphorylated ubiquitin molecule or it can be phosphorylated on its Ubl domain. In the first case, the activation signal (pUb) is fixed and pre-set, while in the second, it depends on the interaction between PINK1 and parkin. Both paths lead to a conformational change in which binding of pUb or pUbl to RING0 releases the catalytic RING2 domain (R2). Upon activation, parkin adds more ubiquitin molecules to the mitochondrial surface, which generates a positive feedback cycle of PINK1 activity and parkin recruitment. Phosphorylation sites are represented by red circles, and the parkin active-site cysteine by yellow sphere when inactive and a star when active.

While the physiological importance of parkin activation by pUb in Parkinson’s disease is unknown, there is a clear effect in cell culture assays of mitochondrial recruitment and mitophagy. Loss of the pUbl-binding site from the K211N mutation has more severe consequences than loss of either the S65 phosphorylation site or the entire Ubl domain (Figure 1). Feedforward activation of parkin may provide a more robust response to PINK1 activity and speed up the positive feedback loop responsible for the switch-like, all-or-nothing behavior observed in cells. A crowded mitochondrial surface with multiple pUb molecules would favor direct feedforward activation; we found poly pUb chains were highly effective in *in vitro* UbVS and autoubiquitination assays of parkin (Figure 4). However, the crowding does not appear to generate contacts between parkin molecules as no complementation between mutants was detected (Figure 1C). The absence of complementation *in vivo* and *in vitro* rules out parkin dimerization as an obligatory step in catalysis (Sauve *et al*, 2018).

Parkin contains two binding sites for phosphoserine with markedly different properties and function. The site on RING1 functions to recruit parkin to pUb on mitochondria. It binds pUb with nanomolar affinity and is extraordinarily selective. pUbl does not compete for binding and, in fact, parkin phosphorylation *increases* the affinity of pUb binding. This is due to allosteric coupling in the RING1 domain between pUb binding and release of the unphosphorylated Ubl domain. The coupling serves to promote parkin phosphorylation and to reduce dissociation of parkin from mitochondria after activation (Kazlauskaite *et al*, 2015; Kumar *et al*, 2015; Sauve *et al*, 2015; Wauer *et al*, 2015a). When phosphorylated, the Ubl domain loses affinity for its site in the autoinhibited conformation. This helps promote its binding to RING0 and increases the affinity of pUb binding.

The second phosphoserine binding site is on RING0 and functions to activate parkin through release of the catalytic RING2 domain (Gladkova *et al*, 2018; Sauve *et al*, 2018). The site was thought to be low affinity and specific for pUbl; however, neither statement is true. The weak binding is a consequence of competition between pUbl and RING2 and does not reflect the true intrinsic affinity of the RING0 site for pUbl. As an intramolecular ligand, the RING2 domain has a high local concentration, and its displacement is energetically costly. Measuring the absolute binding affinity is difficult, but we can deduce that pUbl has higher intrinsic affinity than RING2 since pUbl is able to displace RING2 when both are present in the same polypeptide chain. As a very rough estimate, assuming that autoinhibited parkin is 99% inactive, the local concentration of RING2 is 10 mM, and phosphorylated parkin is 99% active, then the intrinsic affinity for binding pUbl would be 1 µM. Surprisingly, while RING0 site is not strongly selective, it appears to have higher intrinsic affinity for pUb over pUbl. When added *in trans* as free proteins, pUb was between 5 and 10 times more effective than pUbl in activating parkin in functional assays and also showed stronger binding in NMR measurements (Figures 2 & 5).

Several lines of evidence suggest that there is some positive, cooperative coupling between the two phosphoserine binding sites. UbVS assays with the W403A mutant showed slightly more RING2 release when the RING1 site was occupied (Figure 2). Similarly, NMR experiments showed stronger binding of ^15^N-pUbl to the R0RBR/pUb complex than to R0RBR alone. Previous hydrogen-deuterium exchange experiments with phosphorylated parkin detected less protection of RING2 when pUb was present than when it was not (Sauve *et al*, 2018). The largest synergistic effect was manifested in the autoubiquitination assays where pUbl addition to GST-R0RBR in the absence of pUb was relatively ineffective in inducing polyubiquitination (Figure 2E). However, the interpretation is not as clear cut as in the UbVS assays as it is possible that pUb-binding stimulated other aspects of the autoubiquitination reaction such as E2∼Ub binding.

The activation of parkin without its Ubl domain suggests that fusion of the catalytic R0RBR and Ubl domain was a late event in evolution. In primordial parkin, a single pUb-binding site would have sufficed to regulate its recruitment and activation. The subsequent development and specialization of two binding sites - one for recruitment and one for activation - would lead to parkin as we know it today. Distinct sites are necessary to prevent the phosphorylated Ubl domain from interfering with mitochondrial recruitment and so addition of the RING0 domain to the catalytic RBR core must have preceded acquisition of the Ubl domain. Unlike the recruitment site, which must be specific for pUb, there is no reason for the activation site to be specific for pUbl binding and, in fact, the persistence of pUb binding to RING0 argues for a conserved biological function.

The observation of pUb binding to RING0 may also be relevant for understanding other possible functions of parkin in Parkinson’s disease. A variety of results suggest that parkin plays a role in membrane trafficking in nerve terminals. It associates with synaptic membranes in a phosphorylation-dependent manner and can ubiquitinate the synaptic proteins: endophilin and synaptojanin (Cao *et al*, 2014; Trempe *et al*, 2009). Synaptojanin is mutated in an early onset form of Parkinson’s disease and endophilin is a risk locus for Parkinson’s disease (Chang *et al*, 2017; Koros *et al*, 2017). Both are substrates of the Parkinson’s disease associated kinase LRRK2 (Arranz *et al*, 2015; Pan *et al*, 2017). The unexpected plasticity of parkin for binding pUb and pUbl suggests that these or other phosphoproteins may bind the RING1 or RING0 sites to control parkin localization and activation. More broadly, these studies offer hope that it may be possible to find small molecules that modulate parkin for treating Parkinson’s disease.

## Material and methods

### Mitochondria GFP-parkin recruitment time-lapse microscopy

Human osteosarcoma U2OS cells stably expressing a fusion protein of GFP with human parkin (https://www.addgene.org/45875/) were seeded on a 35-mm Glass Bottom 4 Compartment Dish (Greiner Bio-One) (Tang *et al*, 2017). After 18 h, cells were stained with MitoTracker DeepRed FM (ThermoFisher) at a final concentration of 50 nM following manufacturer’s guidelines. Cells were then transferred to a heated stage maintained at 37 °C and 5% CO_2_ using a Zeiss temperature controller and cell perfusion system. Cells were treated with CCCP at a final concentration of 20 µM. Microscopy was performed on a Zeiss AxioObserver.Z1 inverted fluorescent microscope. Fully automated multidimensional acquisition was controlled using the Zen Pro software (Zeiss). Fixed exposure times were as follows: GFP 150 ms; and MitoTracker 40 ms. Images were taken at 2 min intervals for a total of 120 min. Parkin recruitment was visualized by the appearance of punctate GFP fluorescence overlapping Mitotracker fluorescence. The percentage of cells exhibiting Parkin recruitment to the mitochondria was calculated at 2 min intervals over 120 minutes. Approximately 500 cells were examined per mutant over two separate experiments. The fluorescence intensity of GFP-positive puncta was not considered in Parkin recruitment analysis. For statistical analysis, a two-way analysis of variance (ANOVA) with Bonferroni post-test was performed.

### Mitophagy assay

Mitophagy was assessed using flow cytometry analysis of mitochondrially targeted mKeima (a gift from A. Miyawaki, Laboratory for Cell Function and Dynamics, Brain Science Institute, RIKEN, Japan). U2OS cells stably expressing an ecdysone-inducible mt-Keima were induced with 10 µM ponasterone A, transfected with GFP-parkin for 20 h and treated with 20 µM CCCP or an equivalent volume of DMSO for 4 h (Tang *et al*, 2017). Complementation experiments included co-expression of cerulean fluorescent protein (CFP, https://www.addgene.org/54604/) fusions. For flow cytometry analysis, cells were trypsinized, washed and resuspended in PBS prior to analysis on a Thermo Attune NxT cytometer (Thermo) equipped with 405, 488, and 561nm lasers and 610/20, 620/15, 530/30, 525/50 filters (NeuroEDDU Flow Cytometry Facility, McGill University). Measurement of lysosomal mitochondrially targeted mKeima was made using a dual-excitation ratiometric pH measurement where pH 7 was detected using 610/20 filter and excitation at 405nm and pH 4 using 620/15 filter and excitation at 561 nm. For each sample, 100,000 events were collected and single GFP-parkin-positive cells were subsequently gated for mt-Keima. Data were analysed using FlowJo v10.7.2 (Tree Star). The percentage of cells with an increase in the 405:561nm ratio (i.e. cells within the upper gate) was quantified and normalized to the percentage observed in GFP-WT-parkin CCCP treated samples for each replicate. For statistical analysis, a one-way ANOVA with Tukey’s post-test was performed on data from three independent experiments.

### Cloning, expression, and purification of recombinant proteins in *E. coli*

Single-point mutations and deletions were generated using PCR mutagenesis (Agilent). Gibson assembly (NEB) was used to produce Ub-R0RBR chimera: a PCR fragment containing ubiquitin gene and a PCR fragment containing rat parkin residues 77-465 were introduced into the BamHI/XhoI sites of pGEX-6P1 (GE Healthcare). The serine (S77) following glycine 76 of ubiquitin, was mutated in proline to prevent chimera cleavage after ubiquitin glycine 76 by *E*.*coli* endogenous proteins during protein expression. A320R mutation was introduced in the chimera to prevent pUb binding to the high affinity pUb binding site on RING1.

All protein expressions were done in BL21 (DE3) *E. coli* and purification of full-length parkin, R0RBR, Ub-R0RBR, pUb, pUbl, UbcH7, Tc-PINK1 and human His-E1 were done using methods previously described (Berndsen & Wolberger, 2011; Sauve *et al*, 2015). Briefly proteins were purified by glutathione-Sepharose (Cytiva) or Ni-NTA agarose (Qiagen) affinity chromatography, followed by either 3C protease cleavage to remove the GST tag or Ulp protease cleavage to remove the His-SUMO tag. Size-exclusion chromatography was used as a last step. ^15^N-labeled Ubl and Ub were produced in M9 minimal medium supplemented with ^15^NH_4_Cl. Phosphorylated Ub and Ubl were produced and purified according to published procedures (Ordureau *et al*, 2014; Wauer *et al*, 2015a). Purified proteins were verified using SDS-PAGE analysis. Protein concentrations were determined using UV absorbance. Tetra-phospho-ubiquitin chains (pUb)_4_ were obtained from Boston Biochem.

### Ubiquitin vinyl sulfone assays

Ubiquitin vinyl sulfone assays (UbVS) were performed by incubating full-length, chimera Ub-R0RBR or R0RBR, WT or mutant with an excess of UbVS (Boston Biochem) in 25 mM Tris pH 8.5, 200 mM NaCl, 5 mM TCEP for 30 min at 37 °C. Reactions were stopped by addition of 5 x SDS-PAGE loading buffer with 100 mM dithiothreitol (DTT) and analyzed on SDS-PAGE gel stained with Coomassie blue or by western blotting. For western blotting, the samples on SDS-PAGE gel were transferred to Immune-Blot PVDF membrane. The membrane was first blocked with 5% BSA in TBS-T (0.05% Tween 20) overnight at 4 °C and then incubated with 1:2,000 dilution of rabbit anti-Parkin antibody (Ab15954 AbCam) in the blocking solution. The membrane was washed with TBS-T and incubated with 1:10,000 dilution of horseradish peroxidase (HRP)-coupled goat anti-rabbit IgG antibodies (Cell Signaling) in TBS-T. After washing in TBS-T, chemiluminescence was detected using ECL Prime (Cytiva) and bands were visualized with film.

### Ubiquitination assays

The autoubiquitination assays of GST-R0RBR parkin were performed at 22 °C for 5 min by adding 2 µM GST-R0RBR WT, W403A or K211N, to 50 nM human His-E1, 2 µM UbcH7, 100 µM ubiquitin in 50 mM Tris pH 8.0, 150 mM NaCl, 1 mM TCEP, 5 mM ATP and 10 mM MgCl_2_ in the presence of increasing concentrations of pUbl or pUbΔG76. pUbΔG76 was used to prevent incorporation of pUb into the ubiquitination cascade by E1. Reactions were stopped by the addition of 5 × SDS-PAGE loading buffer and the level of ubiquitination was analyzed on SDS-PAGE gels stained with Coomassie blue.

The autoubiquitination assays of full-length parkin and Ub-R0RBR were done in two steps: first, the proteins were phosphorylated by TcPINK1, then the autoubiquitination reaction was carried. The phosphorylation step was performed at 30 °C for 3 hours in 50 mM Tris pH 8.0, 100 mM NaCl, 1 mM TCEP, 0.1% CHAPS, 5 mM ATP, 10 mM MgCl_2_ with 0.1 µM GST-PINK1 and 4 µM parkin. Aliquots were analyzed on Phos-tag SDS-PAGE and quantification of phosphorylation of parkin and Ub-R0RBR was performed by Coomassie Blue band quantifying using the ImageJ software. Autoubiquitination reactions were performed at 22 °C for 1 hr by adding 3.3 µM phosphorylated or non-phosphorylated parkin or Ub-R0RBR to 50 nM human His-E1, 3 µM UbcH7, 75 µM His-ubiquitin S65A in 50 mM Tris pH 8.0, 150 mM NaCl, 1 mM TCEP, 4 mM ATP and 8 mM MgCl_2_. The S65A mutation prevents ubiquitin phosphorylation by any PINK1 present. Reactions were stopped by the addition of 5 × SDS-PAGE loading buffer and the level of ubiquitination was analyzed on SDS-Page gels stained with Coomassie blue.

### NMR experiments

Proteins used for NMR studies were buffer exchanged into 20 mM 4-(2-hydroxyethyl)-1-piperazineethanesulfonic acid (HEPES), 120 mM NaCl, 2 mM DTT, pH 7.4. For pUbl titration, 120 μM unlabeled rat R0RBR/pUb were initially added to 60 μM ^15^N-labeled pUbl. Then, 60 μM and 120 μM of pUb were added to the protein mixture. For pUb titration, 100 μM unlabeled human R0RBR/pUb were added to 50 μM ^15^N-labeled pUb. ^1^H-^15^N correlation spectra were acquired at field strengths of 600 MHz using Bruker spectrometers equipped with a triple-resonance (^1^H, ^13^C, ^15^N) cryoprobe. Spectra were processed using NMRpipe and analyzed with SPARKY (Delaglio *et al*, 1995).

## ACKNOWLEDGMENTS

This work was supported by a Michael J. Fox Foundation grant to K.G., Canadian Institutes of Health Research grants (FDN 154301) to E.A.F and (FDN 159903) to K.G. and Canada Research Chairs (Tier 1) awards to E.A.F and K.G.

## Figures

**Supplemental Figure S1.**
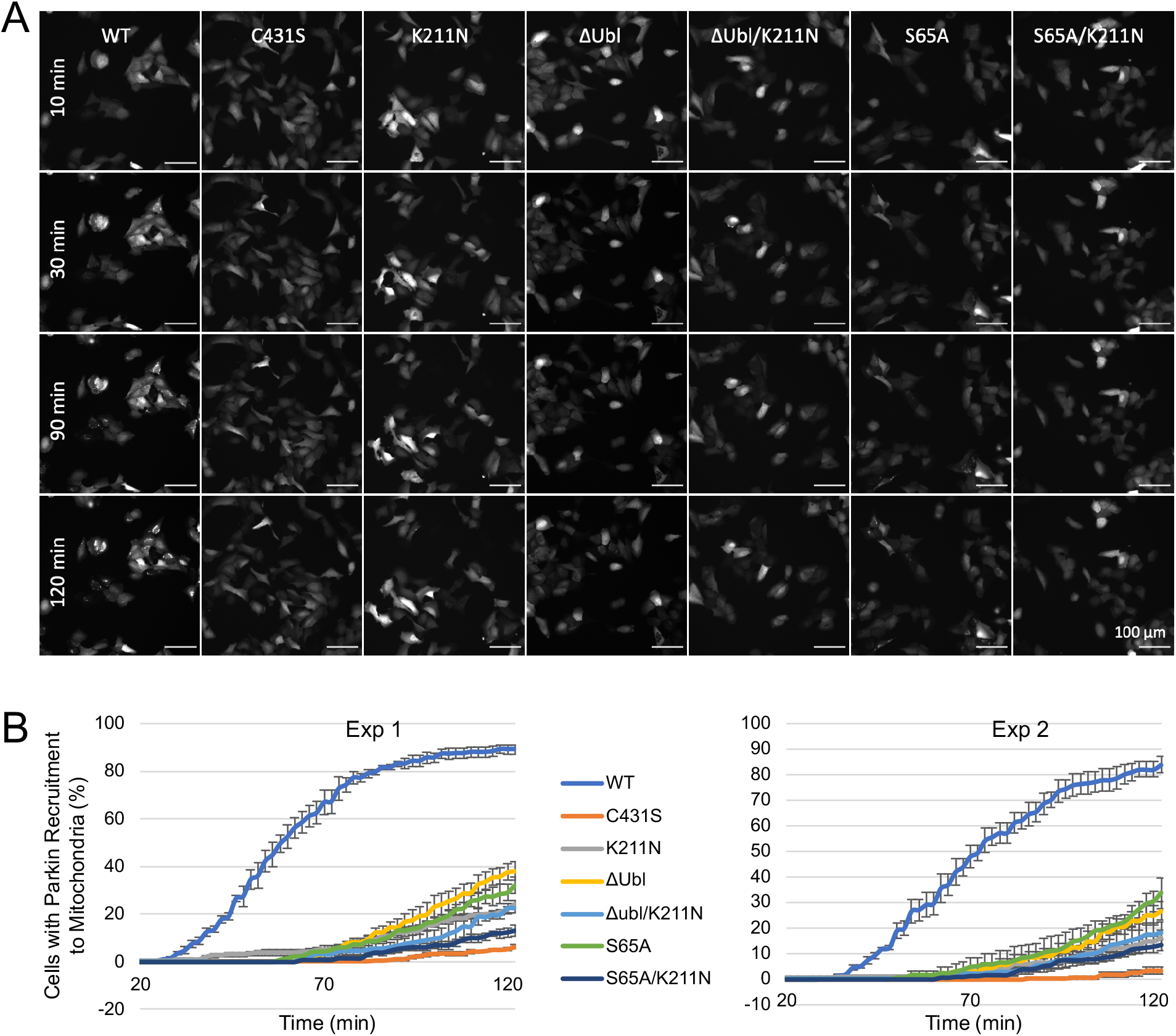
**A**. Time-lapse imaging of parkin recruitment to mitochondria upon treatment with 20 μM CCCP in U2OS cells stably expressing WT, K211N, C431S, ΔUbl (residues 1-76 deleted), ΔUbl K211N, S65A, S65A K211N GFP-parkin. Recruitment can be visualized by the appearance of punctate GFP fluorescence. Scale bar: 100 μm. **B**. Percentage of cells showing parkin puncta determined every 2 min for 120 min after addition of CCCP. Error bars are s.e.m. from four images per time point.

**Supplemental Figure S2.**
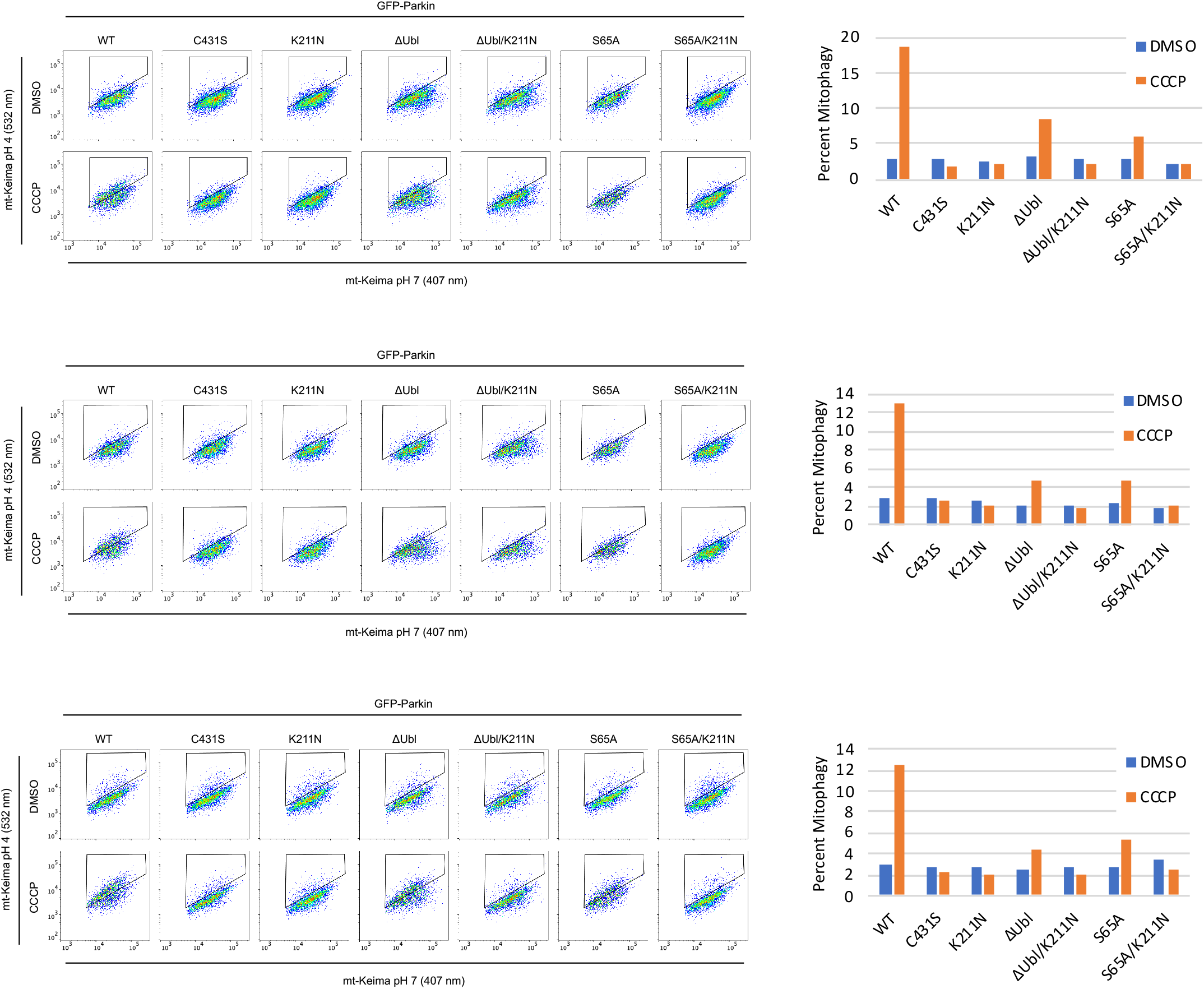
Effects of the loss of the pUbl-phosphorylation site on mitophagy. Three replicates of FACS-based analysis of mt-Keima acidification of U2OS cells transfected with WT, K211N, C431S, ΔUbl, ΔUbl K211N, S65A, S65A K211N GFP-parkin and treated with 20 μM CCCP for 4 hrs.

**Supplemental Figure S3.**
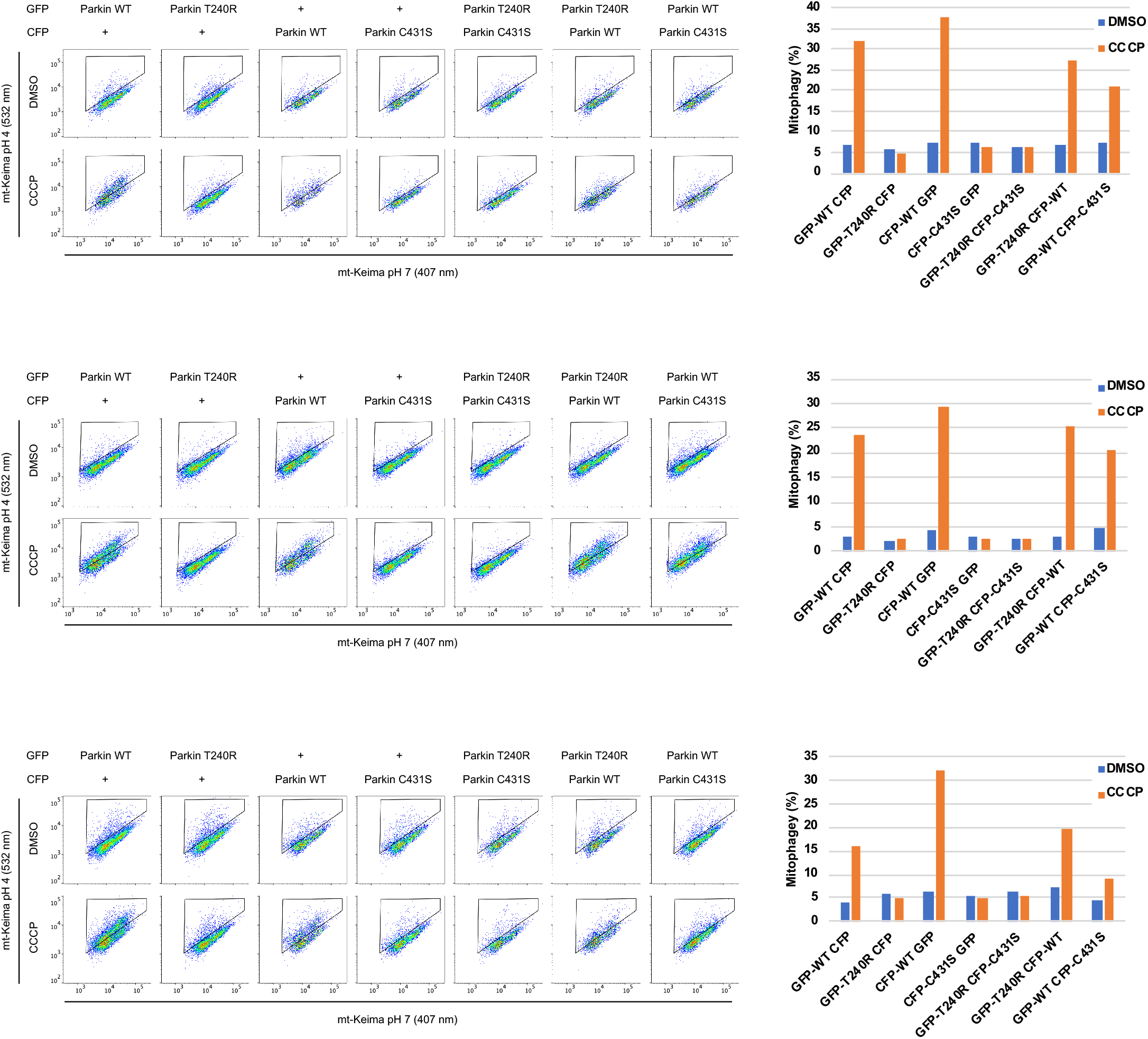
Absence of mutant complementation in mitophagy. Three replicates of FACS-based analysis of mt-Keima acidification of U2OS cells transfected with GFP, CFP, and GFP/CFP-parkin fusion proteins and treated with 20 μM CCCP for 4 hrs.

**Supplemental Figure S4.**
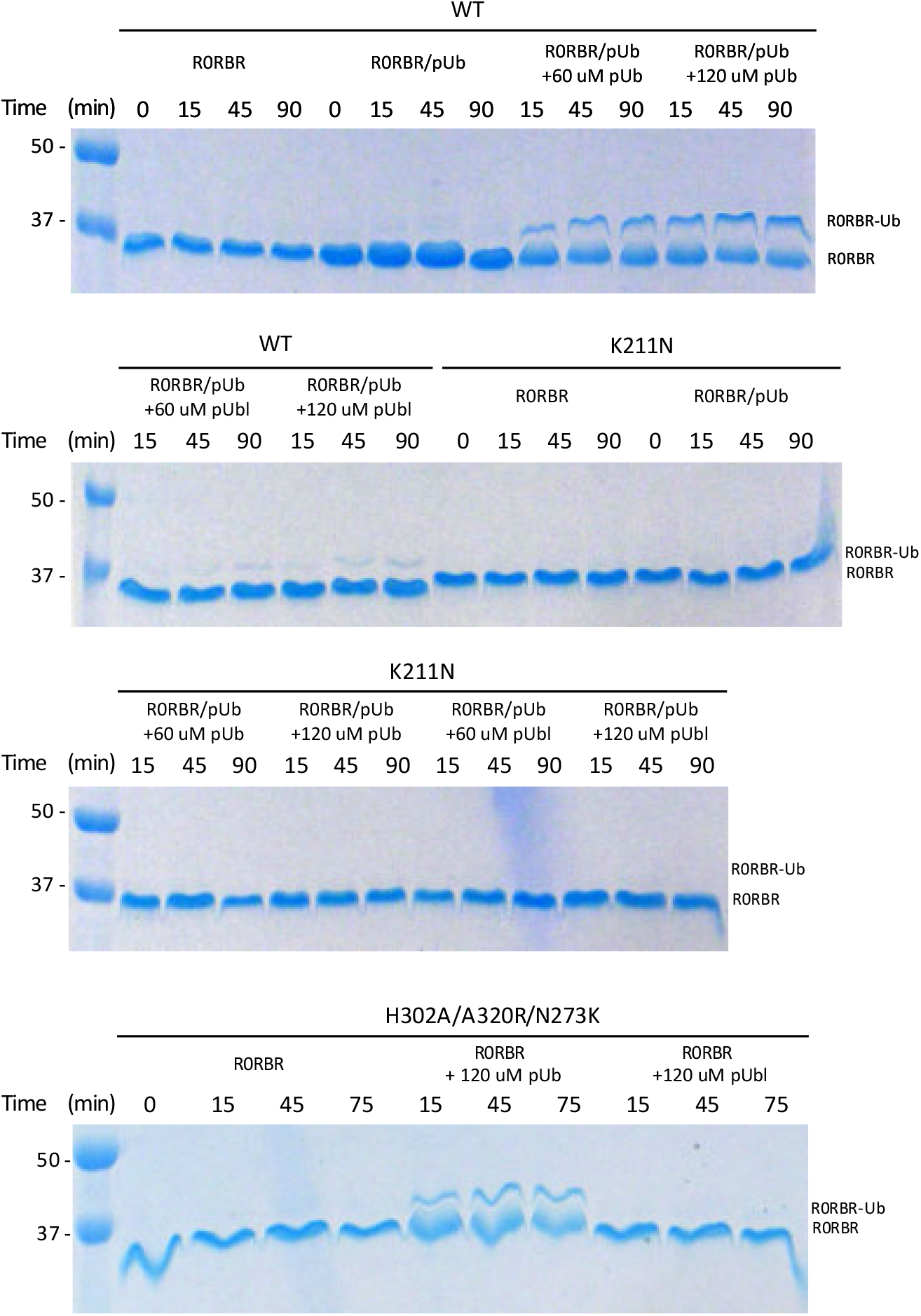
UbVS assays of parkin activation. pUb or pUbl were added to the complex of R0RBR/pUb (WT or K211N) or the R0RBR H302/A320R/N273K prior to addition of 10 µM UbVS. The final R0RBR concentrations were 5 µM. The reactions were performed at 37 °C for a total time of 90 minutes in 25 mM Tris, pH 8.5, 200 mM NaCl, 5 mM TCEP. Reactions were stopped by the addition of 5 × SDS-PAGE loading buffer with DTT (40 mM), and the level of ubiquitination was analyzed on 10% Tris-glycine gels stained with Coomassie blue.

**Supplemental Figure S5.**
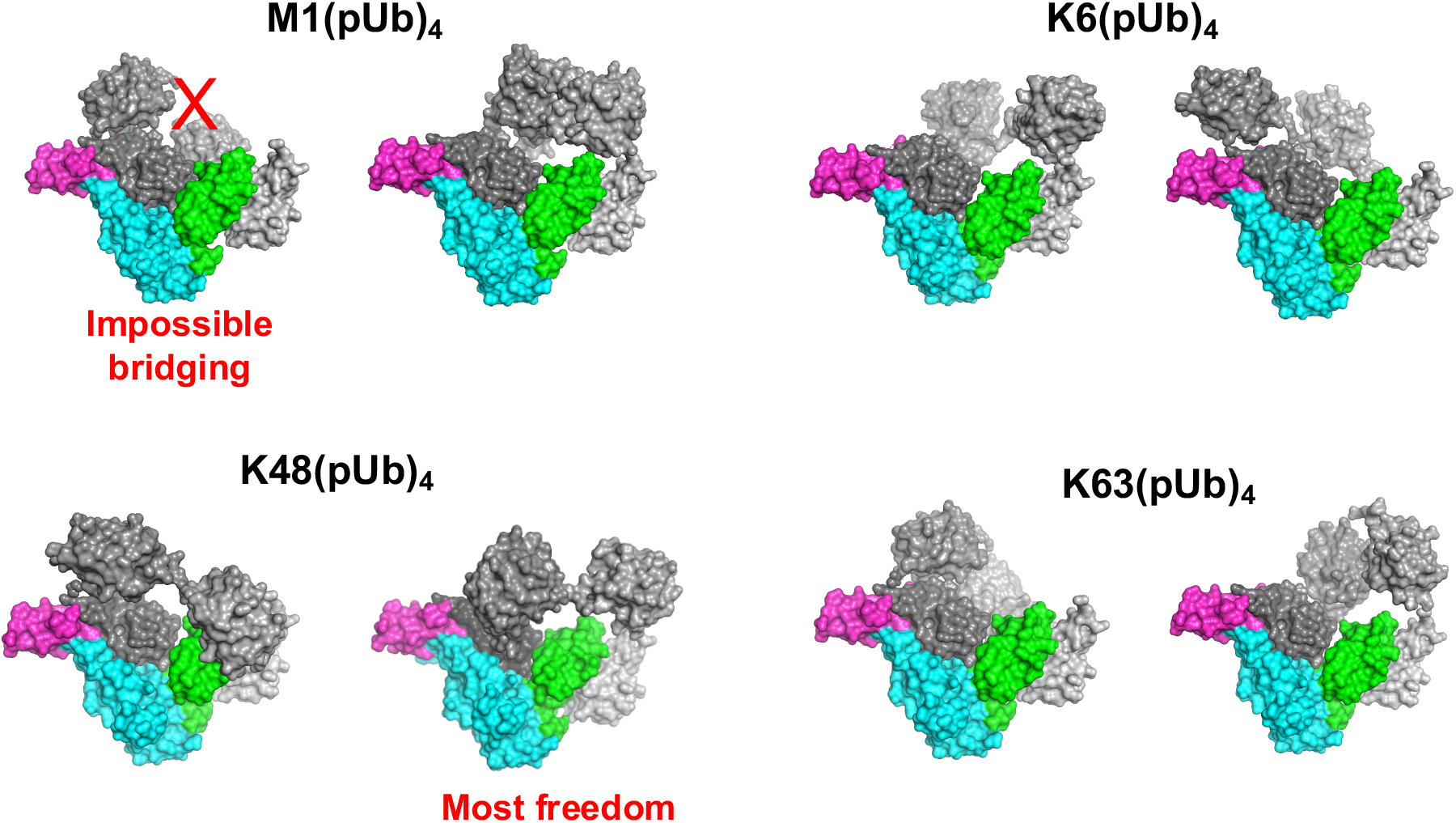
Models of different types of tetra-pUb chains bridging the RING1 and RING0 binding sites of parkin. Parkin RING0 is displayed in green, RING1 in cyan, IBR in magenta, and the pUb chains in grey. Two orientations are shown for each pUb chain.

**Supplemental Figure S6.**
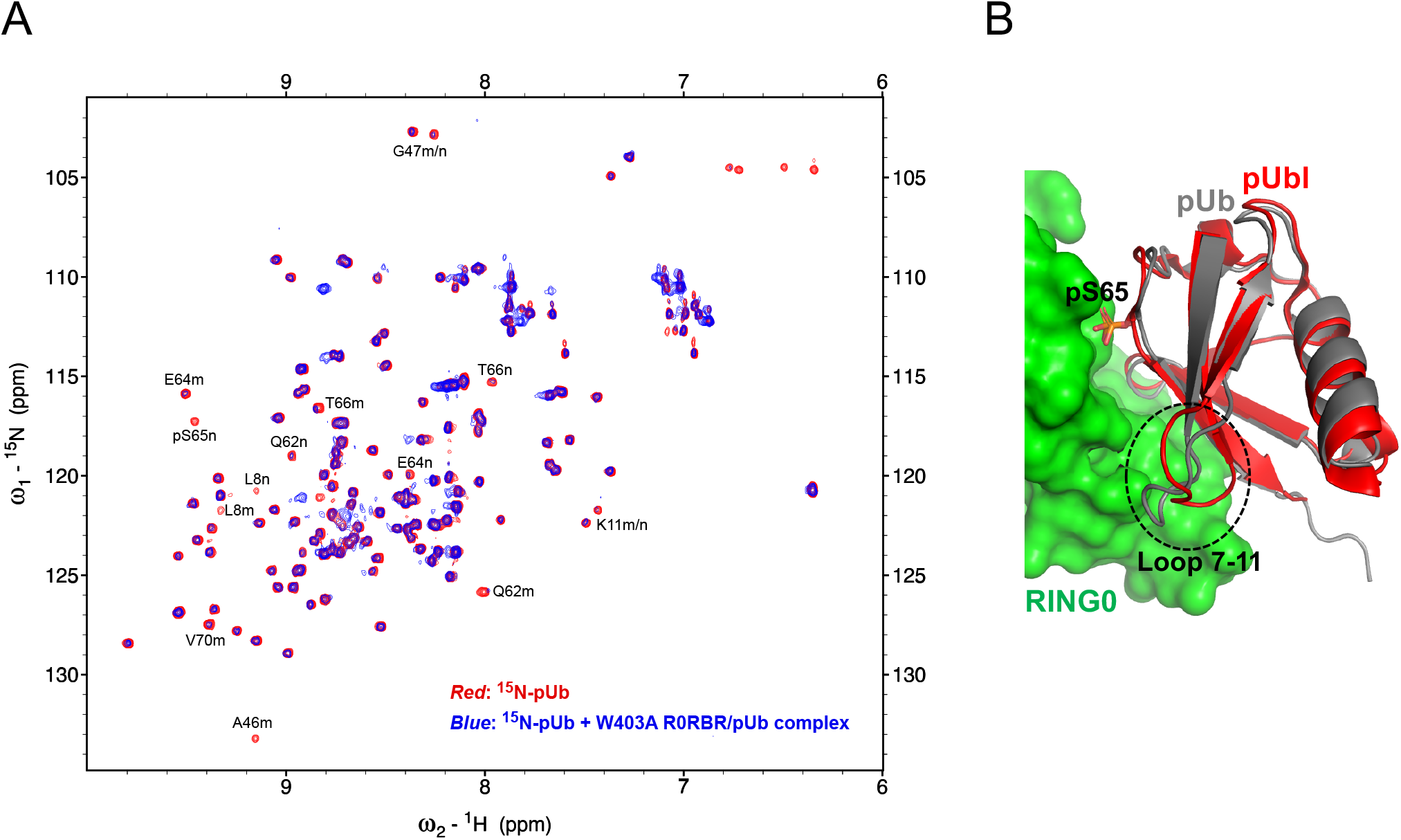
**A**. NMR correlation spectrum (*red*) of ^15^N-pUbl alone (50 µM) overlaid with the spectrum (*blue*) of ^15^N-pUbl (50 µM) in the presence of the parkin W403A R0RBR and pUb complex (100 µM). Signals that are strongly attenuated by binding are labeled by residue number and *m* for the major or *n* for minor pUb conformation. **B**. Model of pUb binding pUbl binding site on RING0. pUb was overlaid over pUbl in pParkinΔREP-RING2-pUb structure (PDB ID 6GLC) (Gladkova *et al*, 2018). RING0 is displayed in green, pUbl in red, and pUb in grey.

